# Graphene-based thin film microelectrode technology for *in vivo* high resolution neural recording and stimulation

**DOI:** 10.1101/2022.11.16.515761

**Authors:** Damià Viana, Steven T. Walston, Xavier Illa, Jaume del Valle, Andrew Hayward, Abbie Dodd, Thomas Loret, Elisabet Prats-Alfonso, Natàlia de la Oliva, Marie Palma, Elena del Corro, Bruno Rodríguez-Meana, María del Pilar Bernicola, Elisa Rodríguez-Lucas, Thomas A. Gener, Jose Manuel de la Cruz, Miguel Torres-Miranda, Fikret Taygun Duvan, Nicola Ria, Justin Sperling, Sara Martí-Sánchez, Maria Chiara Spadaro, Clément Hébert, Eduard Masvidal-Codina, Sinead Savage, Jordi Arbiol, Anton Guimerà-Brunet, M. Victoria Puig, Xavier Navarro, Blaise Yvert, Kostas Kostarelos, Jose A. Garrido

## Abstract

Neuroprosthetic technology aims to restore nervous system functionality in cases of severe damage or degeneration by recording and stimulating the electrical activity of the neural tissue. One of the key factors determining the quality of the neuroprostheses is the electrode material used to establish electrical communication with the neural tissue, which is subject to strict electrical, electrochemical, and mechanical specifications as well as biological and microfabrication compatibility requirements. This work presents a nanoporous graphene-based thin film technology and its engineering to form flexible neural implants. Bench measurements show that the developed microelectrodes offer low impedance and high charge injection capacity throughout millions of pulses. In vivo electrode performance was assessed in rodents both from brain surface and intracortically showing high-fidelity recording performance, while stimulation performance was assessed with an intrafascicular implant that demonstrated low current thresholds and high selectivity for activating subsets of axons within the sciatic nerve. Furthermore, the tissue biocompatibility of the devices was validated by chronic epicortical and intraneural implantation. Overall, this works describes a novel graphene-based thin film microelectrode technology and demonstrates its potential for high-precision neural interfacing in both recording and stimulation applications.

## Introduction

Neural implants offer therapeutic options to patients suffering from certain neurological disorders and other neural impairments (e.g. deafness^1^, Parkinson’s disease,^2^ amputations,^3^ etc.).^4,5^ To further increase the acceptance of neural implants as a therapy, additional improvements in efficacy are needed so that the therapeutic approach outweighs the risks associated with device implantation.^6–10^ Current clinical technology mostly consists of implantable devices that either electrically record or stimulate the nervous system using millimetre-scale metallic electrodes. To improve the interface with the nervous system, the electrode dimensions should be in the micrometre-scale.^3,5,10,12^ By reducing the electrode size to this scale, neural recordings can be captured at a higher spatial resolution, resulting in improved neural decoding.^9,13,14^ Additionally, the reduced electrode size improves neural stimulation focality and expands the gamut of stimulation protocols available to re-create natural neural activation patterns occurring in healthy nervous tissue.^10,15,16^

Due to their unique combination of material properties, graphene and graphene-related materials have emerged as attractive candidate materials for electrode fabrication in chronic bidirectional neural interfaces.^12,17,18^, a monolayer of sp^2^ hybridised carbon atoms, is one of the strongest, more electrically conductive, and stable materials known to date.^19^ In the field of neural interfaces, graphene electrodes offer a capacitive interaction in aqueous media over a wide potential window and mechanical flexibility.^12,20^ Single-layer graphene microelectrodes have been used for neural interfacing applications, but the limited electrochemical performance of this carbon monolayer constrains the potential for miniaturization.^17^ In order to improve performance, multilayer porous electrodes have been explored^18^, although its development has proven to be very challenging due to the stringent requirements of high porosity yet densely packed layers as well as high ion-accessible surface area and low ion transport resistance, which are crucial for the realization of high-performance neural interfacing. Current achievements have lowered the impedance and increased the charge injection limit (CIL) of graphene-based electrodes. However, to date only bulky electrodes of hundreds of micrometres thick have been demonstrated,^21^ which limits the potential integration of the technology into dense arrays for use with anatomically congruent neural interfaces.

In the present work we describe a graphene-based thin film electrode material for neural interfacing. Our Engineered Graphene for Neural Interface (EGNITE) material, leverages the volumetric surface area of nanopores and a high layer packing to increase the electrochemical capacitance by a factor 10^4^ compared to monolayer graphene. Further, we demonstrate a scalable wafer-scale fabrication process of flexible EGNITE microelectrode arrays for high spatial resolution neural recording and stimulation, overcoming the challenges associated with the microelectronic processing of high CIL thin-films. EGNITE microelectrodes exhibit low impedance, high CIL, and current pulse stimulation stability. The fabricated devices have been used *in vivo* to assess their performance for bidirectional neural interfacing in rodents. Cortical and intracortical recording studies have been performed to assess their ability to record spontaneous and evoked local field potentials and multiunit activity and their stimulation performance has been studied using intraneural placement within the rat sciatic nerve to explore the ability to induce selective muscle activation. Additionally, chronic in vivo biocompatibility studies have been performed to assess tissue response. Overall, the results suggest that the EGNITE microelectrode arrays are highly promising for high-precision bidirectional neural interfacing.

## Material Synthesis and Device Fabrication

In this study, we present the synthesis of a micrometer-thick nanoporous graphene-based material (EGNITE) and an extensive structural and chemical characterization. An outline of the fabrication process is shown in **Figure 1a**, and further described in **Methods**. In brief, the EGNITE film is obtained by filtration of an aqueous solution of GO flakes through a porous membrane forming a free-standing GO film that is then transferred on top of the final substrate and hydrothermally reduced to improve the electrical properties and make the material suitable for bidirectional neural interface.

**Figure 1.**
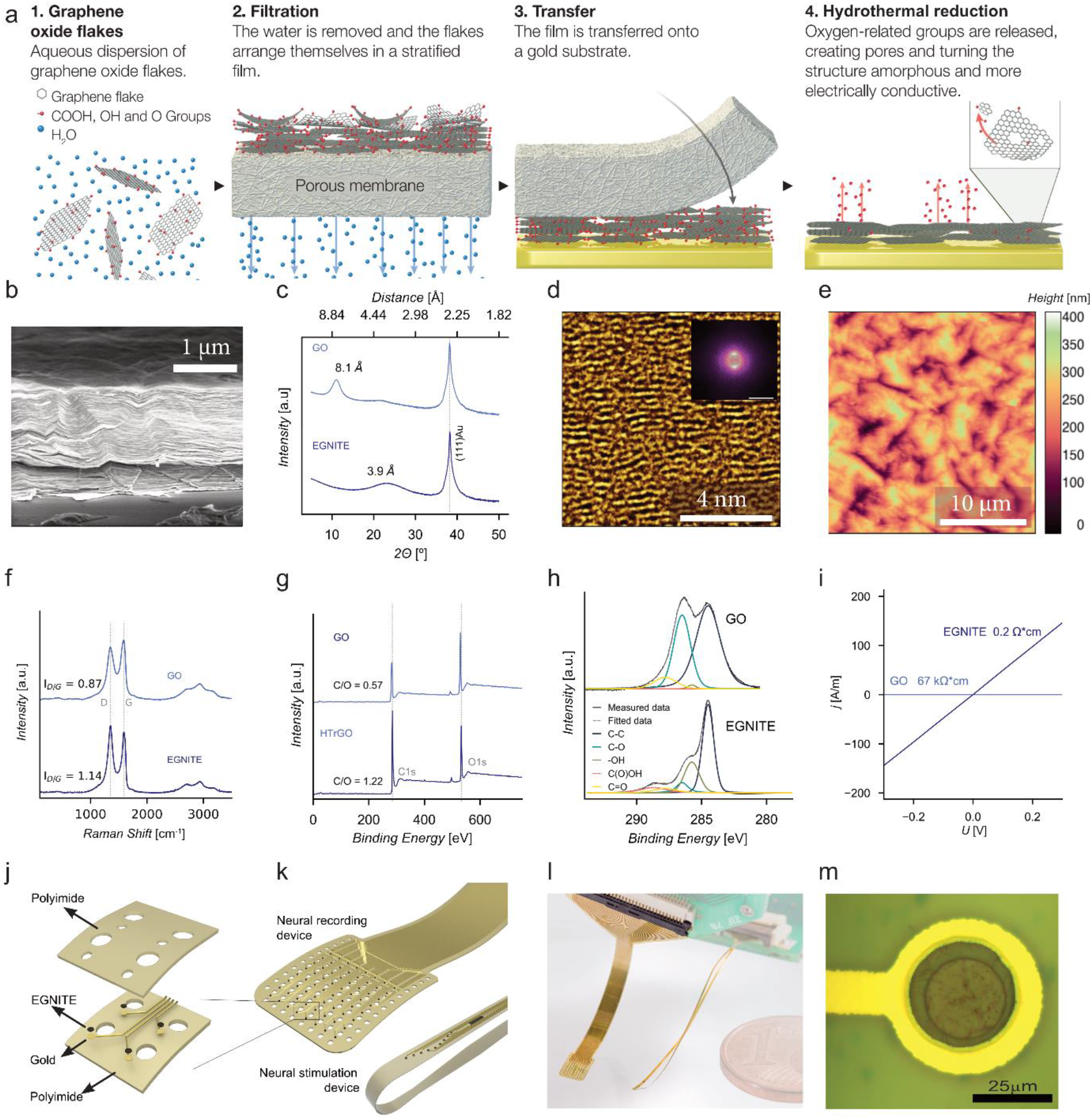
From graphene oxide suspensions to a thin-film electrode: EGNITE technology. **a**. Fabrication process of the graphene thin film, EGNITE. It consists of (1,2) filtering a GO solution through a nitrocellulose membrane, (3) transferring the deposited film of stacked GO flakes onto a conductive substrate and (4) the hydrothermal reduction of the ensemble, which turns the film highly porous and conductive. **b**. SEM micrograph of a cross section of the material. **c**. XRD of GO and EGNITE. The films show a peak corresponding to the parallel stacking. The initial interplanar distance of 8.1 ± 0.8 Å shrinks to 3.9 ± 0.6 *Å* after the hydrothermal treatment. **d**. HRTEM false-colour cross-sectional view of EGNITE and (inset) corresponding power spectrum that shows two symmetric diffuse spots that indicate preferred stacking direction in the material and slight fluctuation of the flakes interplanar distance. **e**. AFM image revealing roughness of the upper surface of the EGNITE film. **f**. Raman spectra of the GO and EGNITE. The ratio between D and G peaks increases after the hydrothermal treatment. **g**. XPS full spectrum and **h**. C1s peak of (top) GO and (bottom) EGNITE. The decrease of the oxygen signal indicates the reduction of the GO film. **i**. Conductivity of the GO and EGNITE films. **j**. Layers involved in the fabrication of flexible arrays of microelectrodes of EGNITE. **k**. Device designs used in this work. **l**. Photography of flexible implants based on EGNITE. **m**. Detailed micrograph of an EGNITE microelectrode of 25 μm diameter.

The structure of the EGNITE film consists of horizontally stacked flakes as revealed by scanning electron microscope (SEM), **Figure 1b**. Following the hydrothermal reduction process, the stacking distance decreases from 8.1 ± 0.8 Å to 3.9 ± 0.6 Å, as assessed by X-ray diffraction (XRD) (**Figure 1c**). The reduction of stacking distance is attributed to the removal of the oxygen groups from the basal plane of the flakes.^22^ The nanostructure cross-section of EGNITE was further investigated by high resolution transmission electron microscopy (HRTEM) (**Figure 1d**). HRTEM data confirmed the stacked configuration of the flakes and the presence of nanometre-scale pores that form capillaries between flake planes. The power spectrum (Fast Fourier Transform) analysis of lower magnification HRTEM images **Fig. SI 1** indicates that the stacked configuration extends across the bulk of the material, with the distance between flakes close to 4 Å, in agreement with the XRD data.

Beyond the internal structure of the material, understanding the topography and atomic composition of the outer surface is of particular importance due to its intended use in direct contact with biological tissue. Atomic force microscope (AFM) measurements shown in **Figure 1e** reveal a surface with a root mean square roughness as low as 55 nm. Further assessment of the film was achieved using Raman spectroscopy. The Raman spectrograms in **Figure 1f** show the G band associated with carbon (1582 cm^-1^), and the D band related to defects in the graphene basal plane (1350 cm^-1^).^23^ The slightly higher defects content observed in EGNITE compared to untreated GO and their nature, associated to vacancies, is tentatively explained by the effect of the hydrothermal reduction of the functionalizing oxygen containing groups, which is assumed to pull out part of the basal plane, creating holes in the reduced GO flakes.^24^ The holes interconnect the capillaries between the flakes planes and are at the origin of the highly enlarged electrochemical surface area of the material, ^24^ as discussed below.

The outer and inner chemical composition of the graphene thin film was studied by X-ray photoelectron spectroscopy (XPS) and electron energy loss spectroscopy (EELS), respectively. **Figure 1g** shows XPS survey spectra which confirms the reduction process during the hydrothermal treatment, decreasing the oxygen content from 25% in the GO film to 12% after the hydrothermal process.^25^ High-resolution XPS measurements in the energy range of C1s peak are shown in **Figure 1h**. The C1s peak is deconvoluted according to the peaks associated to the sp^2^ (284.5 eV) and sp^3^ (285.0 eV) orbitals as well to the ones related to C-OH (285.8 eV), C=O (286.7 eV) and C(O)OH (288.5 eV) bondings.^25^ The analysis shows a strong decrease after the hydrothermal treatment of the C-O peak associated to epoxide groups; but a small contribution of C-OH, C=O and C(O)OH due to hydroxyls, carbonyls and carboxyls remains after reduction. The deconvolution of the O1s peak confirms such behaviour (**2**). The main contribution of the C1s signal after the hydrothermal reduction, however, comes from sp^2^ hybridized C-C orbitals. ^25,26^ The composition deep inside the material was further investigated using EELS in Scanning TEM (STEM) mode, in a FIB cross-section lamella. The cross-sectional study reveals carbon and oxygen relative atomic contents of 85% and 15%, respectively, in the bulk of the material **(Fig. SI 3**, which concurs with the XPS data. According to literature, materials with similar chemical compositions have been reported to be biocompatible.^27^ Regarding the electrical properties, hydrothermal reduction of GO to EGNITE leads to a decrease in the measured resistivity from 67 kΩ·cm to 0.2 Ω·cm (**Figure 1i**), as seen for other hydrothermal treatments.^28^

To exploit EGNITE for neural interfacing, we developed a wafer-scale fabrication process to integrate arrays of EGNITE microelectrodes into flexible devices (**Figure 1j, *Experimental***). Although fabrication of flexible metallic electrode arrays is a standard technology, the integration of a porous micrometer-thick free-standing membrane into microfabrication processing is challenging; so to overcome the integration challenges of EGNITE thin-films we needed to develop a high-yield, large-area transfer of porous GO membrane to the polymer-coated silicon wafer and electrode surface treatments to achieve a strong adherence of the material to the underlaying metal contact. The biocompatible and flexible polymer polyimide (PI) has been used as substrate and insulation,^29^ and Au is used for the tracks. The procedure, further described in **Fig. SI 4**, results in high-yield flexible arrays of EGNITE electrodes that can be fabricated on a wafer scale.

In this study we used an EGNITE film of around 1 μm thickness as a trade-off between electrochemcial performance and array flexibility. Different array designs with 25 μm diameter microelectrodes were fabricated for electrophysiological studies, namely epicortical and intracortical arrays for brain activity recording and stimulation and a transverse intrafascicular multichannel electrode (TIME) for electrical stimulation of peripheral nerves. **Figure 1k (top)** shows the designed 64-channel micro-electrocorticography (μECoG) array organized in an 8×8 grid with a pitch of 300 μm. **Figure 1k (bottom)** shows the fabricated TIME device with two linear arrays of 9 microelectrodes separated by 135 μm designed to selectively stimulate different muscular fascicles inside the sciatic nerve in rodents. The devices have a total thickness of 13 μm and therefore are highly flexible (**Figure 1l**),^20,29^ and contain well-defined 25 μm diameter EGNITE microelectrodes successfully integrated with high-yield (**Figure 1m**).

### Electrochemical characterization

The electrochemical performance of the 25 μm diameter EGNITE microelectrodes was assessed *in vitro* in phosphate buffer saline (PBS) solution (see ***Experimental***). **Figure 2a** shows the cyclic voltammetry (CV) between -0.9 and +0.8 V (vs Ag/AgCl) for EGNITE microelectrodes. The cathodic and anodic charge storage capacitance (cCSC and aCSC) of EGNITE are 33.7 and 27.6 mC/cm^2^, respectively (N=3). **Figure 2b** shows electrochemical impedance spectroscopy (EIS) of EGNITE microelectrodes. At 1 kHz, the EGNITE microelectrodes exhibited an impedance of 26.8 kΩ ± 4.7 kΩ. As a reference, the electrode impedance at 1 kHz for 25 μm diameter electrodes using gold, platinum, PEDOT:PSS/Au, and PEDOT:PSS/Pt electrode materials have been interpolated to be approximately 1.4 MΩ, 1 MΩ, 35 kΩ, and 40 kΩ.^30^ EIS data was fitted with an equivalent electric circuit in which the interface was modelled as a distributed constant phase element (CPE) with a series resistance (Rs).^31^ From the fitting, the interfacial capacitance was found to be 12.54 mF/cm^2^, which corresponds to a 10^4^-fold increase with respect to the typical value of single-layer graphene (2 μF/cm^2^).^32,33^

**Figure 2.**
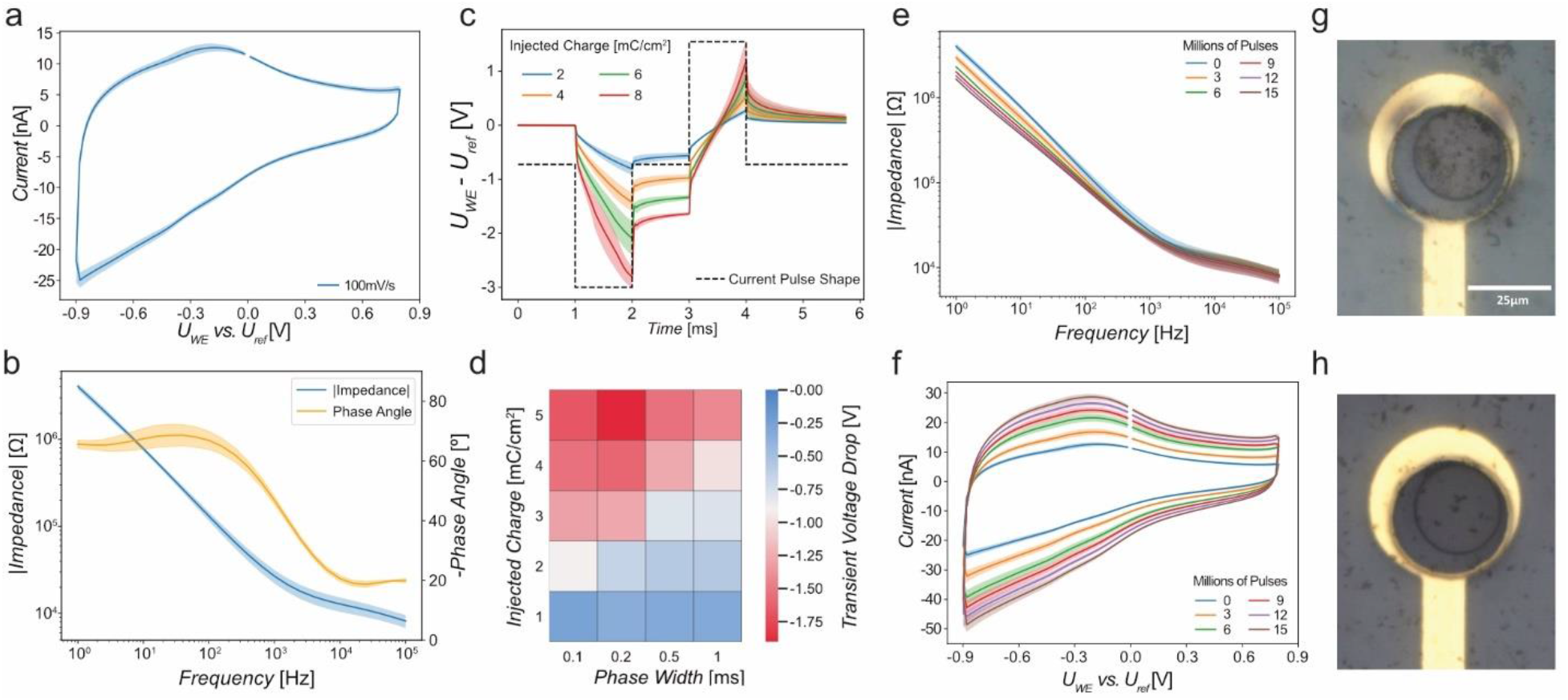
Electrochemical assessment of microelectrodes based on EGNITE. **a**. Cyclic voltammetry at 100 mV/s of EGNITE microelectrodes. **b**. Electrochemical impedance spectroscopy of EGNITE microelectrodes; it shows the module and phase of the impedance vs frequency. **c**. Voltage response to current-controlled biphasic pulses of 1 ms/phase (dashed lined) applied through EGNITE electrodes, corresponding to charge injection values of up to 8 mC/cm^2^. **d**. Map of the cathodic capacitive voltage excursion occurred at the interface between EGNITE microelectrodes and the electrolyte during the injection of current pulses at different levels of injected charge and pulse widths. **e**, Impedance spectra and **f**, Cyclic voltammetry (100 mV/s scan rate) measured throughout continuous stimulation with 15 μA (3 mC/cm^2^) biphasic pulses of 1 ms/phase. Image of 25 μm diameter EGNITE microelectrode before (**g**) and after (**h**) 15 million pulses of continuous stimulation. Data are mean (solid lines) ± SD (shaded area), n = 3 for (**a-c**) and (**e-f**). Data for (**d**) is the mean of the cathodic capacitive voltage excursions of 3 electrodes, n = 1 for each pulse shape.

The performance of EGNITE microelectrodes under current injection has also been studied. **Figure 2c** shows the potential excursion that a 25 μm diameter EGNITE microelectrode experiences upon the injection of rectangular, biphasic current pulses (1 ms/phase) at charge densities of 2, 4, 6, and 8 mC/cm^2^. A quasi-linear voltage shift of maximum 0.9 V exists between the ohmic drops at the edges of the pulses, indicating a capacitive-like behaviour of the electrode-electrolyte interface within the limits of the potential window. EGNITE electrodes have a CIL of 2.8 ± 0.5 mC/cm^2^ for 1-ms pulses. **Figure 2d** shows a map of the voltage polarization experienced by EGNITE microelectrodes in response to biphasic current pulses between 0.1 ms and 1 ms and injected charge densities up to 5 mC/cm^2^. For short pulses of 0.1 ms, approximately 2 mC/cm^2^ could be safely injected since the volumetric capacitance of the electrode could not be fully charged.^11^ As a reference, the charge injection limit for 25 μm diameter electrodes of gold, platinum, PEDOT:PSS/Au, and PEDOT:PSS/Pt electrode materialshave been interpolated to be 0.2, 0.8, 1.9, and 2.7 mC/cm^2^, respectively. ^30^

An assessment of the long-term neural interfacing potential of the microelectrode technology has also been performed. The stability of the electrodes was investigated during continuous electrical stimulation. EGNITE microelectrodes were stimulated with 1 ms biphasic (1 ms interphase gap), 3 mC/cm^2^ pulses at 100 Hz while the impedance and charge storage capacitance (CSC) were monitored every 0.5 million pulses. EGNITE microelectrodes (N=3) were observed to be stable after 15 million pulses as indicated by the rather constant impedance at 1 kHz (**Figure 2e**). However, a decrease of the impedance at lower frequencies is observed during stimulation; the fitted capacitance shows an increase from 12.5 mF/cm2 to 34.4 mF/cm2. In accordance with the impedance data, the average CSC increases from 61.7 to 136.7 mC/cm2 throughout the stimulation (Figure 2f) and the CIL increases up to 3.3 ±0.2 mC/cm2. We tentatively attribute the capacitive increase when pulsing to a slow increase of the total electrochemical surface area due to the porous nature of the EGNITE electrodes. No obvious structural changes were observed in the electrode as illustrated in **Figure 2g,h**. The high performance of the EGNITE electrodes is attributed to the combination of the capacitive interaction in aqueous media with the large electrochemical surface area obtained during the hydrothermal reduction.

Furthermore, we investigated the mechanical stability of EGNITE electrodes by sonication of the devices while immersed in an ultrasound water bath. After consecutive sonication for 15 min, 200 W and 15 min, 300 W the EGNITE electrodes remained attached to the device **Fig. SI 5**. No delamination nor cracking of the electrodes was observed.

### Recording neural signals

#### Epicortical Device implantation & recording

The suitability of the EGNITE microelectrode technology to record *in vivo* neural signals was assessed by using flexible micro-electrocorticography (μECoG) devices (**Figure 1k**, top) to monitor the neural activity at the surface of the auditory cortex of anesthetised Sprague Dawley rats (4-months old) **(Figure 3a**). The μECoG devices contain an array of 64 EGNITE microelectrodes (Ø 25 μm, pitch 300 μm). The probe was positioned over the left auditory cortex (AI and AAF regions) (**Figure 3b**). The average impedance of the microelectrodes was 58 ± 25 kΩ at 1 kHz **Fig. SI 6**. Pure tones of 16 kHz and 200 ms duration (3 ms rise and 20 ms fall times) were presented at 90 dB SPL to generate evoked activity. **Figure 3c** shows the 10 Hz high-pass filtered signal from each of the 64 EGNITE electrodes in the array over a 350 ms window in which a pure tone was presented. The large negative and positive voltage peaks that are observed after the onset and offset of the sound stimulus are evoked local field potentials (eLFP).^**34**^

**Figure 3.**
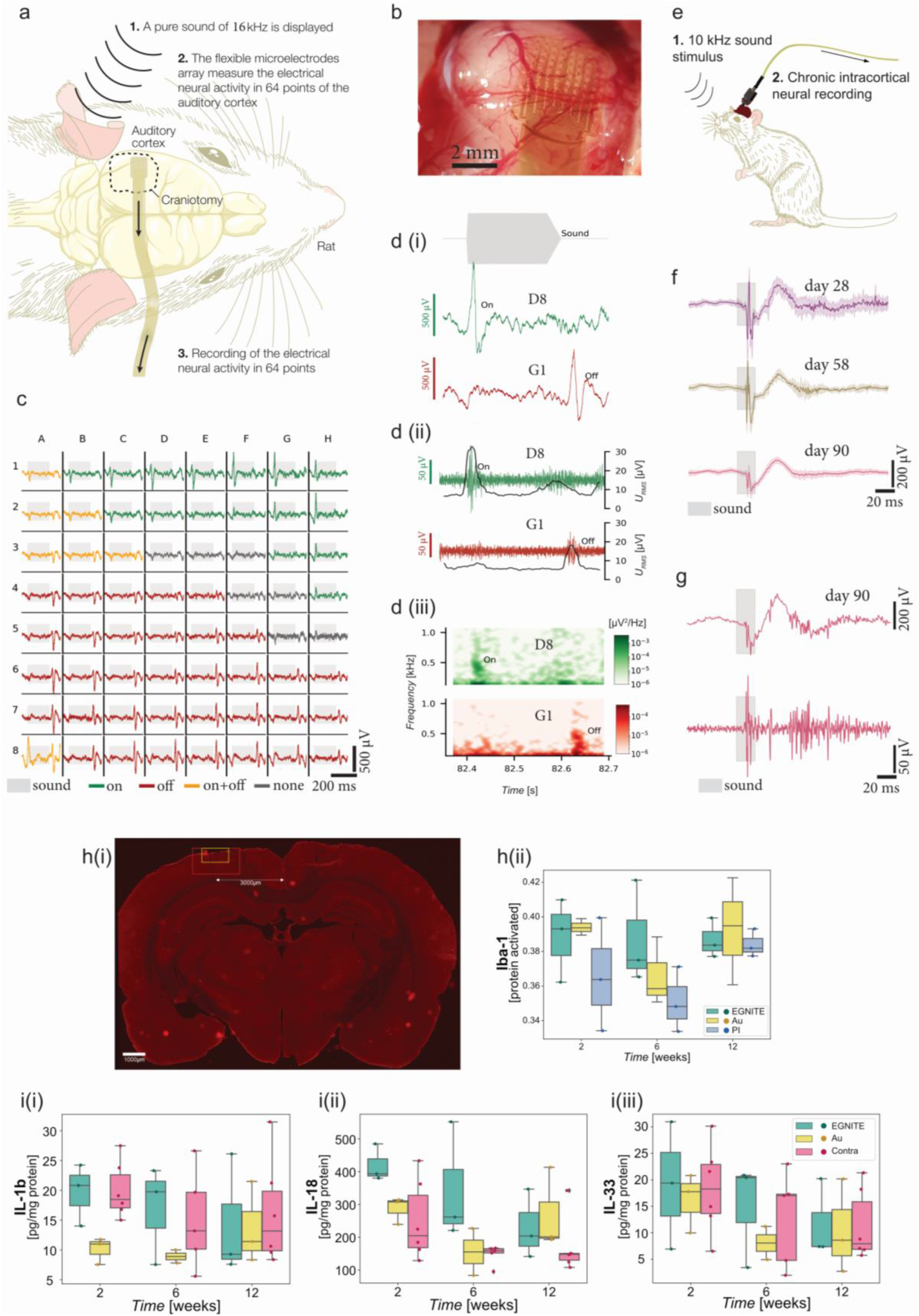
EGNITE-based microelectrode arrays map with high spatiotemporal fidelity local field potentials and MUA for over 90 days. **Recording: a**. Schematic diagram of the acute experiment using an EGNITE μECoG flexible array to record epicortical neural activity of a rat. Evoked activity was induced by a pure 16-kHz tone stimulus. **b**. EGNITE μECoG array on the auditory cortex of a rat. **c**. Mapping of the evoked neural activity depicting a single event across all 64 EGNITE microelectrodes. Depending on the region, onset (green), offset (red), both (yellow), or no onset/offset (dark grey) responses are recorded. **d (i)**. Response of regions G1 (onset, green) and D8 (offset, red) to the evoked local field potentials. **d (ii)**. These signals are high-pass filtered at 200 Hz to reveal high frequency MUA activity (green and red), confirmed by a simultaneous increase of the RMS value of the signal (grey) **d (iii)**. Spectrogram of the neural activity measured at G1 and D8 during activation and deactivation of the auditory signal. **Intracortical: e**. Schematic diagram of intracortical flexible array configuration to record neural activity about 1.7 mm deep into the prefrontal cortex with EGNITE electrodes. **f**. Averaged auditory evoked potentials recorded 30, 60 and 90 days after the implantation. Data are mean (solid line) ± SD (shaded areas), n = 30. **g**. Single auditory evoked potential high-pass filtered above 10 Hz (top) and above 200 Hz (bottom) at recorded on the 90^th^ day of implantation. **Biocompatibility: h**. Microglial activation in the cortical region in direct contact with the probe was determined based on their phenotype morphology assessment by immunohistochemistry and algorithmic image processing. h(i) Representative brain section, post-implantation (left hemisphere) immunohistochemically stained for Iba-1. Area of interest was set at 3 mm from midline, based on average center of device implantation coordinates. Red box is 8x magnification and was used as input data for image analysis of microglial phenotype morphological analysis. Yellow box is higher (20x magnification). h(ii) Iba-1 signal based on image processing of immunohistochemical sections. No significant differences in Iba-1 expression were found between different implanted device material or timepoints post-implantation. **i**. ELISA-based quantification shown, of **i**(i) anti- and **i**(ii), **i**(iii) pro-inflammatory cytokines present in the brain tissue samples in direct contact with the EGNITE microelectrodes for 2-, 6- and 12-weeks following their epicortical implantation. Cytokine levels were normalised to total protein content (mg/mL) of the tissue sample. Data are mean ± SD, n = 3 for each device material, and between n = 6-9 for contralateral hemispheres. *P < 0.05 (Two-way ANOVA with Tukey’s multiple comparisons test).

Similar mappings were obtained on 2 more animals, as shown in **Fig. SI 6**. The high quality of the recorded signal allows mapping of single-event eLFPs as displayed in **Figure 3c**, which reveals a spatial clustering with ON responses (green) located on the top right corner and OFF responses (red) grouped in the bottom region. The spatial distribution of ON and OFF responses recorded during single events can vary from event to event; **Fig. SI 7** shows a selection of single event recordings together with the average response of the array to the 16 kHz tone stimulation. **Figure 3d(i)** shows details of the ON and OFF single trial signals recorded in the two main clusters, extracted from regions G1 and D8 of the device (as indicated in **Figure 3c**), respectively. The amplitudes of the evoked response range between 500 μV and 700 μV and have latencies, of 30 ms and 34ms after the onset of the sound stimulus, for the ON and OFF stimulus respectively, in good agreement with previous reports. ^34^ **Figure 3d(ii)** shows the same signals from regions G1 and D8 high-pass filtered at 200 Hz overlapped with its root mean square (RMS) value. The high frequency multi-unit activity (MUA) and the corresponding increase of the RMS amplitude due to MUA nicely correlate to the eLFP, indicating a synchronous behaviour of the neurons to the stimulus. ^34^ The spectrograms of the electrical recordings of G1 and D8 (**Figure 3d(iii)**) confirm the increase of high frequency neural activity at the stimulation onset and offset.

The recording capability of EGNITE microelectrodes is further confirmed in recordings of spontaneous cortical activity **Fig. SI 8**. The amplitude of the recorded spontaneous activity exceeds that of the evoked signal due to the contribution of the signals below 10 Hz. The high-pass filtering of the signal at 200 Hz shown in **Fig. SI 8a** uncovers the contribution of low-amplitude (20-30 μV peak-peak), high frequency components of the signal, which are found to disappear post-mortem. The calculation of the RMS noise of the signals allows correlation of the low and high frequency components of the spontaneous activity. The intrinsic RMS noise of the electrode (calculated from the post-mortem recordings) was of 2.5 μV **Fig. SI 8b**, which was very close to the technical capabilities of the electronic setup.^9,35^ **Fig. SI 8c** shows the averaged power spectral density (PSD) obtained from 60 EGNITE electrodes over a 30-minute *in vivo* recording, compared to the PSD of a post-mortem recording. The SNR (calculated from the division of the signals) reaches 40 dB at 10 Hz and 5 dB at 1 kHz (**Fig. SI 8d)**, demonstrating an excellent capability to record high fidelity signals at both low and high frequencies.^36^

To benchmark the recording capabilities, similar measurements were performed using commercially available Pt electrodes (ø 60 μm, NeuroNexus). The average electrode impedance was 695 ± 361 kΩ, which is one order of magnitude larger than the 25 μm diameter EGNITE microelectrodes. Evoked and spontaneous signals of lower amplitude were recorded with Pt electrodes and the calculated SNR was 25 dB at 10 Hz and 2 dB at 1 kHz (see **Fig. SI 9**). The 60μm diameter Pt microelectrodes, despite having an area 5.7 times larger than the 25 μm diameter EGNITE electrodes, were outperformed by a factor of 2 in terms of SNR by the EGNITE microelectrodes.

#### Intracortical Device Implantation and Recording

Additionally, chronic recording experiments (see ***Experimental***) were performed using an EGNITE intracortical device implanted in the prefrontal cortex in a mouse for over 90 days (**Figure 3e**). After the first month, auditory evoked potentials (AEPs) were recorded longitudinally over the three months while the animal was freely moving. **Figure 3f** shows an overlap of 20 averaged trials on days 28, 58, and 90. The observed AEP peak at 40 ms is detected at all timepoints At day 28, AEP is 140 μV, while at day 90 it goes to 90 μV (**Fig. SI 10**), which is consistent with previous reports of increased electrode impedance throughout the first 100 days of intracortical implantation.^37–39^ High-pass filtering of the AEP signal at 10 and 200 Hz reveals that action potentials are detected in vivo in chronic recordings (**Figure 3g**). The quality of the recordings could be demonstrated by the detection of action potentials from individual neurons (**Fig. SI 10**).

#### Cortical Biocompatibility

To assess the biocompatibility of the thin-film EGNITE devices, we carried out chronic (12-week) biocompatibility studies with epicortical implants. The epicortical devices were implanted on the right hemisphere of adult, male Sprague-Dawley rats (**Figure 3h(i)** and **Fig. SI 11**) for up to 12 weeks before histological and immunohistochemical evaluation of microglia phenotype morphology or extraction of cortical brain tissue for cytokine expression (by ELISA) (**Figure 3h,i**). Gold and Polyimide (PI)-based implants were used to compare with the EGNITE devices at three timepoints (2, 6, and 12 weeks (*n* = 3 per group)) post-implantation. Both immunohistochemical and cytokine analysis of the implanted hemisphere was compared to the contralateral hemisphere (without device implantation).

Immunohistochemical analysis by Iba-1 (microglia marker) staining of the brain tissue after device implantation was performed to assess the level of microglial cell activation in the region directly in contact with the implanted devices (**Figure 3h(i)**). The microscopy data were imaged processed using an algorithm for microglia phenotype characterisation. These results indicated no significant differences in the levels of microglia activation, regardless of device material or timepoint (**Figure 3h(ii)**). Cortical tissue samples in contact with the implanted devices were collected from a separate group of animals to assess the level of neuro-inflammation (by ELISA) potentially triggered by the chronic implantation of EGNITE devices. No significant changes in the levels of inflammatory cytokines from the cortical tissue samples at each timepoint (or between the different device types) were found (**Figure 3i** and **Fig. SI1**). A minor increase in IL-1b, IL-6 and MCP-1 can be seen in the 2- and 6-week EGNITE samples compared to those implanted with Gold or PI, however this difference is not seen at the 12-week time point and was not found to be significant (**Figure 3i(i), Fig. SI 11**, respectively). When compared to the contralateral hemisphere at each timepoint, again no appreciable differences could be seen for any of the cytokines quantified, with the exception of the anti-inflammatory marker IL-18 (**Figure 3i(ii)**) which was significantly increased at 2 weeks following implantation with EGNITE compared to the contralateral hemisphere. However, by 12 weeks this reaction also abated to background levels.

### Stimulation of nerve fibers

#### Intrafascicular Device Implantation and Stimulation

To evaluate the stimulation capability of the EGNITE microelectrodes, transverse intrafascicular multichannel electrode (TIME)^40^ devices were implanted transversally in the sciatic nerve of anesthetised Sprague Dawley rats in acute experiments (**Figure 4a**). The device consisted of 2 linear arrays (A and B) of 9 electrodes (ø 25 μm) along a 1.2 mm stripe. Each linear array faced opposite sides of the stripe. Once implanted (**Figure 4b**), the device crossed the peroneal fascicle (responsible for the innervation of the tibialis anterior (TA) muscle) and the tibial fascicle (responsible for the innervation of both the gastrocnemius (GM) and plantar interosseous (PL) muscles). Each electrode in the EGNITE array was individually stimulated and the elicited compound muscle action potentials (CMAP) of the TA, GM, and PL muscles were simultaneously recorded by monopolar needles inserted in the muscles^40^ (**Figure 4c**).

**Figure 4.**
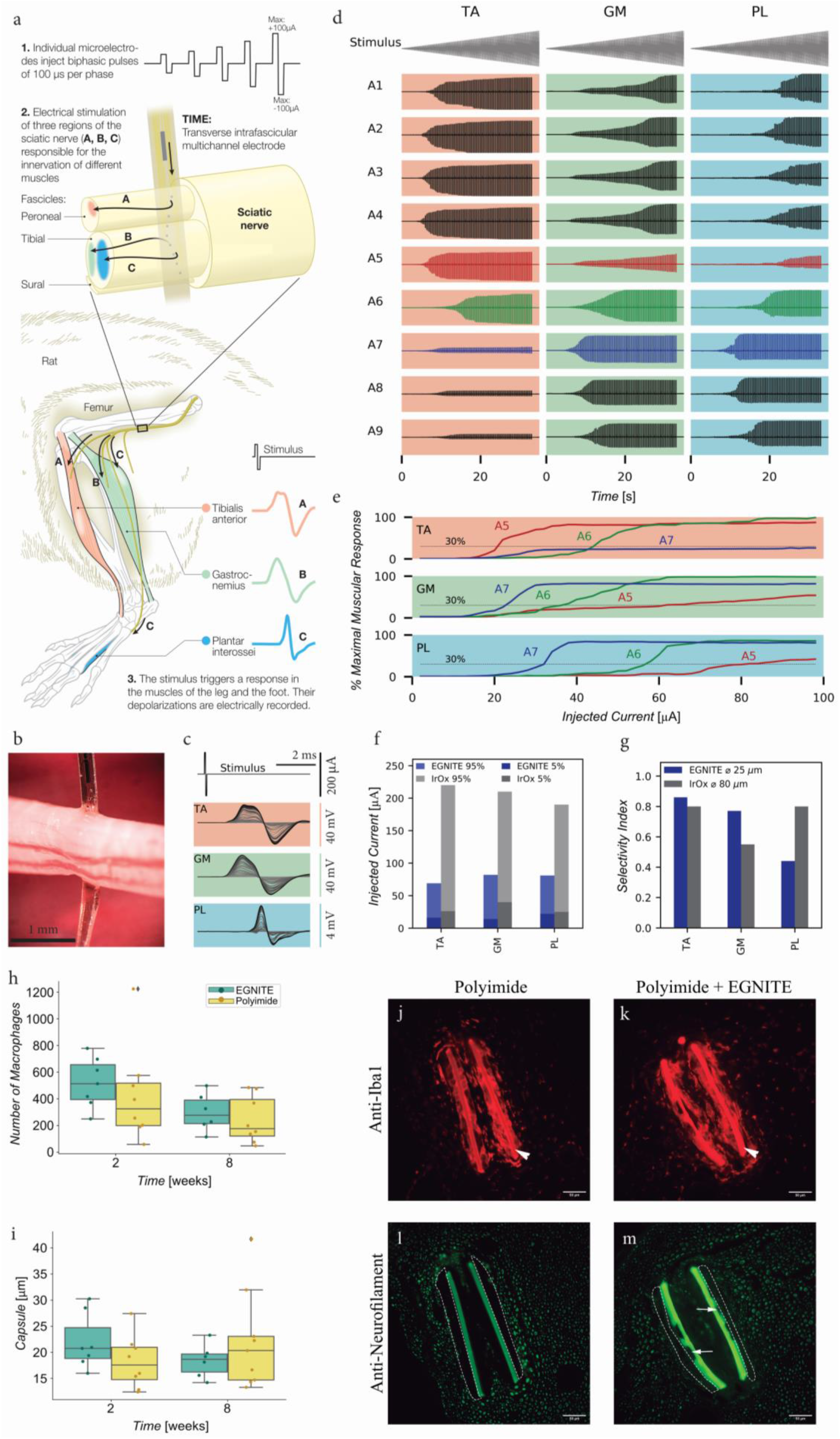
*In vivo* neural stimulation of peripheral nerve and biocompatibility. **Stimulation: a**. Schematic diagram of the acute stimulation experiment. A transversal intra-fascicular microelectrode array (TIME) is implanted in the sciatic nerve of a rat crossing the peroneal and the tibial fascicles. The innervation of TA muscle can be found in the peroneal fascicle, whereas the GM and PL *muscles* innervation is located in the tibial fascicle. Biphasic pulses of 100 μs/phase and a maximum amplitude of 100 μA were injected individually at the 9 microelectrodes of 25 μm diameter located in the device. The local electrical stimulation can depolarize the nerve fibers from its vicinity and trigger the electrical activity of the muscles, which was recorded with needle electrodes. **b**. Optical micrograph of the implanted TIME device in the sciatic nerve. **c**. CMAPs recorded from the TA, GM, and PL muscles in response to increasing levels of injected current pulses applied to one of the electrodes. **d**. Recorded CMAP in TA, GM, and PL muscles in response to trains of biphasic current pulses of increasing amplitude applied to the 9 microelectrodes of the array, on one side of the TIME. **e**. Normalized CMAP of TA, GM, and PL muscles in response to pulses injected to microelectrodes A5-A7 from the implanted TIME. **f**. Comparative plot of the injected current needed to elicit a 5% and 95% of the maximum CMAP using microelectrodes for EGNITE (blue) and IrOx (grey).^43^ **g**. Comparative plot of the selectivity index at the minimal functionally relevant muscular stimulation for EGNITE (blue) and IrOx (grey). ^43^ **Biocompatibility: h**. Plot of the number of inflammatory Iba1+ cells in the tibial nerve after 2 and 8 weeks of EGNITE device implantation. **i**. Tissue capsule thickness around the biocompatibility devices. Data are mean ± SD, n = 7 for each device material. *P < 0.05 (Two-way ANOVA followed by Bonferroni post hoc test for differences between groups or times). **j, m**. Representative images of transverse sections of a tibial nerve at 8 weeks after implantation of biocompatibility devices, made of polyimide alone (j, l) or polyimide with EGNITE (k,m), stained for inflammatory cells (antibody against Iba1, j, k) and for axons (antibody against neurofilament 200, **l, m**). The arrowhead points to the transverse sections of the polyimide strips that were longitudinally inserted in the nerve, that have strong autofluorescence. The thin arrow points to a site with EGNITE in the polyimide strip in m. The tissue capsule is delineated as dotted lines in l and m). Scale bar: 50 μm.

To stimulate the nerve, a train of 100 biphasic pulses (100 μs/phase) with increasing current amplitude (0 to 100 μA, in 1 μA steps) were injected through the microelectrodes at 3 Hz for 33 seconds. **Figure 4d** shows the recorded response of TA, GM and PL muscles to these current pulses applied through the A1-A9 microelectrodes. The recorded signals are normalized to the maximum amplitude of the CMAP. Currents as low as 15-20 μA elicited CMAPs that increased in amplitude until a maximum activation was reached when current was further increased. The obtained recruitment curves reflect the typical sigmoidal shape (**Figure 4e**).^40,41^ The pattern of muscular activation changed depending on the stimulating electrode. The activity could be split in 2 clusters: one for A1-A5, in which the TA muscle was activated at lower stimulus intensity than the GM and the PL muscles, and another one for A7-A9, in which the GM and PL muscles exhibited more activity than the TA muscle. This suggests that the first five microelectrodes were placed in the peroneal fascicle, whereas the last three were located within the tibial fascicle. From these responses, critical parameters for implants aiming at restoring mobility such as current thresholds and selectivity index of the modulated the muscular activity can be derived.^3^

Of particular interest are electrodes A5-A7, which are located nearest the interface between the two peroneal and tibial fascicles. In these electrodes, significant changes of current threshold and selectivity index were observed. Using 30% of muscle activation (the minimum stipulated to overcome gravity^40^) as a benchmark, the current stimulation at which this occurred for each muscle was determined for the TA, GM, and PL muscle as marked in **Figure 4e**. For example, the injected current amplitude needed to evoke TA muscle activity to 30% of maximum was 21 μA when stimulated by electrode A5. From these data, a selectivity index above 0.85 was calculated for the 30% activation of the TA, 0.77 for the GM, while for the PL it was only 0.44. As an indicator, an optimal selectivity index of 1 indicates that one muscle can be solely activated without any activation in the other recorded muscles. ^3,10^ Taking into account that the TA innervation is conveyed through the peroneal fascicle and the innervations of the GM and the PL are contained in the tibial fascicle,^42^ the lower activation thresholds and higher selectivity indices are obtained for stimuli coming through electrodes placed in closer positions, highlighting the importance of small microelectrode sizes.^40,43^ The lower selectivity index obtained by the PL is probably due to the high dispersion of the nerve fibers innervating this muscle within the tibial fascicle compared to the more confined fibers of the TA and GM.^42^

**Figure 4f** shows the minimum current thresholds to achieve a 5% and a 95% of the CMAP amplitude in TA, GM and PL muscles taking into account the activity triggered by all the microelectrodes shown in **Figure 4d**. To reach 5% contraction, less than 50 μA were required for all the muscles, whereas less than 100 μA were needed to reach 95% of maximal activity.

Compared to previous studies in which TIMES arrays with 80μm diameter electrodes of iridium oxide were used,^43^ EGNITE electrodes elicited a response with current thresholds 2 to 3 times lower. The selectivity index values achieved with EGNITE electrodes also improved with respect to those previously obtained using larger IrOx electrodes, indicating higher focality in the stimulation (**Figure 4g**).^10,42,43^ The lower response thresholds and improved selectivity index values are likely a result of using smaller EGNITE electrodes that focally stimulate the highly confined TA and GM nerve fibers at higher charge densities. Thus, the high charge injection limit of EGNITE allows the fabrication of smaller electrodes which can help improve neural stimulation capabilities and eventually therapeutic outcomes. Apart from the therapeutic advantage this can provide, the power consumed by the stimulation device can be significantly reduced and extending its functional lifetime.^3,43^

#### Peripheral Nerve Biocompatibility

To study the biocompatibility of thin film EGNITE devices in sciatic nerve, an intraneural device was designed in which the area of EGNITE in contact with the nerve was increased by a factor of 20, aiming to maximize the contact area of the material with the tissue and to investigate immune responses. These intraneural implants, with and without EGNITE, were longitudinally implanted in the tibial branch of the sciatic nerve of Sprague-Dawley rats (**Fig. SI 10e**,**f**) as in previous studies.^**44**^ Electrophysiological, pain and locomotion functional studies as well as immunohistochemical labelling of the nerve were conducted at 2- and 8-weeks post-implantation and compared to the contralateral nerve or paw (n = 6-8 per group) (see ***Experimental*** and **Fig. SI 10**).

The results of nerve conduction tests did not show any difference in the CMAP amplitude and latency between the implanted paw with polyimide alone, with polyimide plus EGNITE or the contralateral paw during the 8 weeks follow-up (**Fig. SI 11a**,**b**), indicating that there was no damage to myelinated motor nerve fibers by any of the implants. The mechanical algesimetry test, typically used to assess pain, showed no changes between groups at any time point (**Fig. SI 11d**), indicating that there was no damage of small nerve fibers or irritation induced by nerve compression or axonal injury. Finally, the walking track test to assess locomotor function did not show variations between the two types of devices along follow-up (**Fig. SI 11c)**, confirming that there was no functional nerve damage in the rats assessed.

The histological analysis indicated that there were no signs of nerve damage or axonal degeneration induced by the implants. One of the main events during the foreign body reaction (FBR) is the infiltration by hematogenous macrophages into the implanted site, as part of the inflammatory phase.^45^ Comparison between implants with and without EGNITE revealed no differences in the amount of macrophages in the nerve (**Figure 4h, j, k**). The last phase of the FBR and one the main obstacles for long-term functionality of intraneural electrodes is the formation of a fibrous capsule around the implant. **Figure 4i** shows that the capsule thickness formed around the polyimide strips was similar for implants with and without EGNITE at both 2 and 8 weeks, indicating that EGNITE does not induce damage to the nerve and fibrotic scar formation. Immunohistochemical images (**Figure 4j-m**) show numerous axons near the implants (at around 20 μm) of polyimide without and with EGNITE, indicating limited damage and remodelling after implant, consistent with previous works.^44^ Altogether, the chronic biocompatibility study indicates that EGNITE is suitable for chronic intraneural implantation, inducing no significant nerve damage nor neuroinflammatory response.

## Conclusions

In order to exploit the full potential of electronic neural interfaces, there is a need for robust, physiologically compatible materials that allow high-resolution and stable bidirectional neural interfacing. With their combination of structural, electrical, and electrochemical properties EGNITE electrodes are well suited for neural interfacing, based on its highly porous structure (with a 10^4^ increase in capacitance compared to planar graphene electrodes) and a reversible charge injection mechanism. We have demonstrated that the graphene-based EGNITE electrodes designed here can be miniaturized to the microscale and integrated in flexible, wafer-scale, thin-film fabrication neurotechnology while keeping their properties for bidirectional neural interfacing. The developed microelectrodes arrays can sustain long-term stimulation (15 million pulses) at 3 mC/cm^2^ and are mechanically. *In vivo*, the EGNITE microelectrodes exhibited extremely low intrinsic noise levels, in the range of the lowest reported, allowing high SNR during epicortical recordings for mapping brain signals with high spatial resolution. The developed technology can also sustain implantation with stable recordings for up to 3 months.^9,35^ Additionally, during acute *in vivo* stimulation studies conducted in the sciatic nerve, the small diameter (25 μm) and high density of EGNITE electrodes permitted focal stimulation that increased the selectivity and lowered the charge stimulation thresholds required for muscle activation, when compared to recent reports.^43^ This is of particular interest for providing precise motor activation to users of bionic hand prostheses,^4^ and for high resolution deep brain stimulators (DBS), as it is expected that smaller electrodes improve the localization of stimuli to the therapeutic targets.^15,16,46^ Biocompatibility assessments conducted in the cortex and peripheral nerve suggest that the EGNITE material is well tolerated by the local immune response and hence can be used as implant. The high-resolution recording capabilities of EGNITE could also facilitate the discovery of new neural biomarkers, and in combination with selective stimulation this technology paves the way for the development of adaptive closed-loop therapies.^13,14^ Additionally, the low charge stimulation threshold obtained with intrafascicular EGNITE microelectrodes will be beneficial to lower power consumption of chronically implanted neural stimulators and extend their functional lifetime. While further investigation will be necessary to evaluate the chronic functionality and stability of the electrode technology while interfacing with specific systems targeted for neuromodulation therapy,the combination of electrical stimulation and neural recording capabilities, coupled with *in vivo* performance and immune tolerance indicates that the thin-film EGNITE microelectrode technology is a strong candidate for applications requiring bidirectional neural interfacing and enabling closed-loop functionality.^47^

## Acknowledgements

This work has been funded by the European Union Horizon 2020 Research and Innovation programme under Grant Agreement No 881603 (Graphene Core3); FLAG-ERA JTC 2017 project GRAFIN, from the Ministerio de Ciencia, Innovación y Universidades of Spain PCI2018-093029, PCI2018-092935, FLAG-ERA JTC 2021 project RESCUEGRAPH, from the Agencia Estatal de Investigación of Spain PCI2021-122075; FIS2017-85787-R (co-financed by Agencia Estatal de Investigación y el Fondo Europeo de Desarrollo Regional); TERCEL (RD12/0019/0011) and CIBERNED (CB06/05/1105) funds from the Instituto de Salud Carlos III of Spain; and by Fundación Ramón Areces (CIVP18A3897). ICN2 is supported by the Severo Ochoa Centres of Excellence programme, funded by the Spanish Research Agency (AEI, grant no. SEV-2017-0706), and by the CERCA Programme / Generalitat de Catalunya. The co-authors D.V. and S.M-S are supported by the International PhD Programme La Caixa - Severo Ochoa (Programa Internacional de Becas “la Caixa”-Severo Ochoa). D.V. acknowledges that this work has been done in the framework of the PhD in Electrical and Telecommunication Engineering at the Universitat Autònoma de Barcelona. J.dV and E.dC. acknowledge the Ministerio de Economía, Industria y Competitividad of Spain for Juan de la Cierva incorporation fellowships. M.C.S. received funding from the European Union Horizon 2020 under the Marie Sklodowska-Curie grant agreement No 754510 (PROBIST). This work has made use of the Spanish ICTS Network MICRONANOFABS partially supported by MICINN and the ICTS ‘NANBIOSIS’, more specifically by the Micro-NanoTechnology Unit of the CIBER in Bioengineering, Biomaterials and Nanomedicine (CIBER-BBN) at the IMB-CNM. Part of the present work has been performed in the framework of Universitat Autònoma de Barcelona Materials Science PhD program and Neuroscience PhD program. The FIB sample preparation was conducted in the Laboratorio de Microscopias Avanzadas at Instituto de Nanociencia de Aragon-Universidad de Zaragoza. Authors acknowledge the LMA-INA for offering access to their instruments and expertise. This study was also financed by grant PID2019-104683RB-I00 funded by MCIN/AEI/ 10.13039/501100011033 to M.V.P.

The authors would like to acknowledge Mr Pau Nebot for technical assistance. The authors would also like to thank Drs C.Bullock and C.Bussy for their contributions at the early stages of the project. The University of Manchester Bioimaging and Single Cell Genomics Facility microscopes used in this study were purchased with grants from the UKRI Biotechnology and Biological Sciences Research Council (BBSRC), the Wellcome Trust, and the University of Manchester Strategic Fund. The RT97 antibody was obtained from the Developmental Studies Hybridoma Bank developed under the auspices of the NICHD and maintained by the University of Iowa, Department of Biology.

## Author contributions

D.V. did most of the design, fabrication and characterization of the microelectrode arrays, contributed to the design and performance of the in vivo experiments, analysed the data and wrote the manuscript. S.T.W. supported the design, fabrication and characterization of the devices, and participated in the discussion of the analysed data. X.I. designed the neural arrays and supported their fabrication. J.dV. coordinated and performed the in vivo stimulation and supported B.R-M. with the chronic biocompatibility experiments, both contributed to the data analysis. E.P. contributed developing the fabrication process of the neural probes. N.dlO. contributed performing the in vivo stimulation experiments and their data analysis. M.P. contributed performing the in vivo recording experiments. E.dC. performed the AFM and Raman spectroscopy measurements and contributed to their data analysis. B R-M. supported the in vivo stimulation experiments. M.dPB. supported electrochemical characterization of electrodes. E.R-L. coordinated in vivo intracortical experiments. T.A.G. coordinated in vivo intracortical experiments. J.M.dC. contributed developing the fabrication process of the neural probes. M.T.M. supported the fabrication of the devices. J.S. significantly reviewed the document. M.C.M. performed the TEM measurements and contributed analysing the data. S.M.S. and J.A. supported the analysis of the TEM data. C.H. supported the development the EGNITE. E.M.C. wrote and extensively reviewed the manuscript. A.G.B. developed a custom python library and participated in the neural data analysis. M.V.P. designed intracortical recording experiments. X.N. designed the in vivo neural stimulation and chronic biocompatibility experiments and reviewed the manuscript. B.Y. designed and performed the neural recording experiments and reviewed the manuscript. K.K. designed the biocompatibility assessment experiment and reviewed the manuscript. J.A.G. participated in the development of the technology, analysis of the arrays characterization and the neural data. K.K. and J.A.G. conceived, initiated and overall supervised the entirety of this work. All authors read and reviewed the manuscript.

## Conflicts of Interest

D.V., A.G., K.K. and J.A.G. would like to declare that they hold interest in INBRAIN Neuroelectronics that has licensed the technology described in this manuscript.

## Experimental

### Material preparation and characterization

Aqueous GO solution was diluted in deionised water to obtain a 0.15 mg/mL solution and vacuum filtered through a nitrocellulose membrane with pores of 0.025 μm, forming a thin film of GO. The thin film was then transferred to the target substrate and the GO film - substrate stack was hydrothermally reduced to form EGNITE. The base-substrate for all characterization studies of EGNITE was a square (1×1 cm^2^) of Si/SiO^2^ (400 μm / 1μm). **XPS**. XPS measurements were performed with a Phoibos 150 analyzer (SPECS GmbH, Berlin, Germany) in ultra-high vacuum conditions (base pressure 5·10^−10^ mbar) with a monochromatic aluminium Kα x-ray source (1486.74 eV). Overview spectra were acquired with a pass energy of 50 eV and step size of 1 eV and high-resolution spectra were acquired with pass energy of 20 eV and step size of 0.05 eV. The overall resolution in those last conditions is 0.58 eV, as determined by measuring the FWHM of the Ag 3d5/2 peak of sputtered silver. **XRD**. XRD measurements (θ-2θ scan) were performed in a Materials Research Diffractometer (MRD) from Malvern PANalytical. This diffractometer has a horizontal omega-2theta goniometer (320 mm radius) in a four-circle geometry and it worked with a ceramic X-ray tube with Cu *K*_*α*_ anode (λ=1.540598 Å). The detector used is a Pixcel which is a fast X-ray detector based on Medipix2 technology.

#### Raman spectroscopy

Raman spectroscopy measurements were performed using a Witec spectrograph equipped with a 488 nm laser excitation line. For the measurements, Raman spectra were acquired using a 50x objective and the 600 g nm-1 grating; laser power was kept below 1.5 mW to avoid sample heating. **TEM**. A FIB lamella was prepared Helios NanoLab DualBeam (LMA-INA, Zaragoza) for the cross-section study of the EGNITE sample. Structural analyses were performed by means of Transmission Electron Microscopy using a TECNAI F20 TEM operated at 200 kV, including HRTEM and HAADF-STEM techniques. The STEM-EELS experiment was performed in a Tecnai F20 microscope working at 200 KeV, with 5mm aperture, 30 mm camera length, convergence angle 12.7 mrad and collection angle 87.6 mrad. As we used 0.5 eV/px and 250 eV as starting energy in the core-loss acquisition, we did not acquire Si k-edge expected at 1839 eV, the Pt M-edge at 2122 eV and the Au M-edge at 2206 eV. The relative C – O atomic composition has been obtained focusing our attention in the GO layer and assuming that the edges analysed (C and O in our case) sum to 100%. This assumption is valid in our case as evidenced in the SI maps. The energy differential cross section has been computed using the Hartree-Slater model and the background using a power low model. **Electrical conductivity**. Electrical conductivity measurements were performed using a Keithley 2400 sourcemeter in 2-point configuration. The samples measured consisted of EGNITE films of 1×1 cm^2^ on top of a SiO_2_ substrate.

#### Data analysis

XRD, Raman and XPS data were analysed using Python 3.7 packages (Numpy, Pandas, Scipy, Xrdtools, Lmfit, Rampy, Peakutils, Matplotlib). The distance between planes was calculate from XRD measurements according to the Snell’s law. Once the data was moved into the spatial domain, the maximum of the peaks were fitted. The corresponding distance gave a mean value of the distance between planes. Deviations from those mean values were calculated from the FWHM of Lorentzian fittings of the peaks on the spatial domain. XPS and Raman spectroscopy measurements were analysed by fitting a convolution of peaks on expected locations for the corresponding features. The conductivity values of the GO and EGNITE were obtained by fitting the I-V curves measured in the electrical conductivity measurements to the Ohm’s law. Data are n = 1 for each measurement.

#### Flexible array fabrication

Devices were fabricated on 4” Si/SiO_2_ (400 μm / 1 μm) wafers. First, a 10 μm thick layer of polyimide (PI-2611, HD MicroSystems) was spin coated on the wafer and baked in an atmosphere rich in nitrogen at 350 ºC for 30 minutes. Metallic traces were patterned using optical lithography of the image reversal photoresist (AZ5214, Microchemicals GmbH, Germany). Electron-beam evaporation was used to deposit 20 nm of Ti and 200 of Au. After transferring of the GO film, the film was then structured using a 100 nm thick aluminium mask. Columns of aluminum were e-beam evaporated and defined on top of the future microelectrodes via lift off by using a negative photoresist (nLOF 2070, Microchemicals GmbH, Germany). Next, the GO film was etched everywhere apart from the future microelectrodes using an oxygen reactive ion etching (RIE) for 5 minutes at 500 W. The protecting Al columns were subsequently etched with a diluted solution of phosphoric and nitric acids. After, a 3 μm thick layer of PI-2611 was deposited onto the wafer and baked as done before. PI-2611 openings on the microelectrode were then defined using a positive thick photoresist (AZ9260, Microchemicals GmbH, Germany) that acted as a mask for a subsequent oxygen RIE. Later, the devices were patterned on the PI layer using again AZ9260 photoresist and RIE. The photoresist layer was then removed in acetone and the wafer cleaned in isopropyl alcohol and dried out. Finally, the devices were peeled off from the wafer, placed in sterilization pouches and hydrothermally reduced at 134 ºC in a standard autoclave for 3 hours.

### Microelectrode electrochemical characterization

Electrochemical characterization of the microelectrodes was performed with a Metrohm Autolab PGSTAT128N potentiostat in 1X PBS (Sigma-Aldrich, P4417) containing 10mM phosphate buffer, 137 mM NaCl, and <u>2.7 mM</u> KCl at pH 7.4 and using a three-electrodes configuration. An Ag/AgCl electrode (FlexRef, WPI) was used as reference and a Pt wire (Alfa Aesar, 45093) was used as counter.

Prior to performance evaluation, electrodes were pulsed with 10,000 charge-balanced pulses (1 ms, 15 μA). Exposing electrodes to continuous pulsing protocols were proceeded by 100 cyclic voltammetry cycles (−0.9 to +0.8 V) at 50 mV/s, and 20 repetitions of 5000 1-ms pulses and re-determination of open circuit potential.

#### Data analysis

Electrochemical characterization data was analysed using Python 3.7 packages (Numpy, Pandas, Scipy, Pyeis, Lmfit, Matplotlib). Impedance spectroscopy data was fitted to an equivalent electric model consisting of a resistance (R) in series with a constant phase element (CPE). From there, the CPE value was approximated to a capacitance and divided by the microelectrode geometrical area to obtain an equivalent value for the interfacial capacitance of EGNITE. Microelectrode charge storage capacitance (CSC) was calculated from cyclic voltammetry measurements by integrating the cathodic and anodic regimes of the measured current and normalizing by the scan rate. Microelectrode charge injection capacity (CIC) was established by determining the current pulse amplitude that elicited a capacitive voltage polarization that exceeded the electrode electrochemical water window.

#### Statistical analysis

Data are mean ± SD, n = 3 for CV, EIS and chronopotentiometries. Data of the map of cathodic capacitive voltage excursion is the mean of the cathodic capacitive voltage excursions of 3 electrodes, n = 1 for each pulse shape.

### Mechanical stability evaluation – Ultrasound sonication

EGNITE electrode arrays were placed inside a beaker filled with water in an ultrasound water bath (Elmasonic P 180 H). Sonication was applied at 37 kHz, for 15 minutes at 200 W and followed by an additional 15 minutes of sonication at 37 kHz with the power elevated to 300 W. Images of electrodes were acquired before and after the sonication steps.

### Neural Recording: epicortical implantation

All experimental procedures were performed in accordance with the recommendations of the European Community Council and French legislation for care and use of laboratory animals. The protocols were approved by the Grenoble ethical committee (ComEth) and authorized by the French ministry (number 04815.02). Sprague Dawley rats, weighing ∼600 g, were anesthetized intramuscularly with Ketamine (50 mg/kg) and Xylazine (10 mg/kg), and then fixed to a stereotaxic holder. Removing the temporal skull exposed the auditory cortex. Dura mater was preserved not to damage the cortical tissue. A hole was drilled at the vertex in order to insert the reference electrode and a second hole, 7 mm toward the front from the first one, in order to insert the ground electrode. The electrodes were 0.5-mm-thick pins used for integrated circuit sockets. They were placed to make electrical contacts with the dura mater and fixed to the skull with dental cement. We then mounted the surface microelectrode ribbon on the auditory cortex as **Figure 3b** shows. The vein patterns identify the auditory cortex, in area 41 of Krieg’s rat brain map. Cortical signals were simultaneously amplified with a gain of 1000 and filtered with a band-pass of 20 Hz–1500 Hz, 12 dB/octave, and digitized at a sampling rate of 20 kHz. A speaker 20 cm in front of a rats’ ear, contra-lateral to the exposed cortex, delivered acoustic stimuli. The stimuli delivered were monitored by a 1/4-in microphone (Brüel & Kjaer, 4939) placed near the ear and presented in sound pressure level (dB SPL re 20 Pa). We examine the vertex-positive (negative up) MLRs, evoked by alternating clicks at 80-dB SPL, and tone burst stimuli at 70-dB SPL with the frequencies ranging 5 kHz–40 kHz, the rise and fall time of 5 ms and the duration of 200 ms.

#### Data analysis

Electrophysiological data was analysed using Python 3.7 packages (Numpy, pandas, scipy, Neo, Elephant, Sklearn Matplotlib) and the custom library PhyREC (https://github.com/aguimera/PhyREC). RMS values were calculated with a sliding window of 20 ms at frequencies above the 200 Hz. Spectrograms were calculated on a range between 70 Hz and 1.1kHz. PSD was calculated over 60 s of continuous recordings. For a given electrode array, two PSD were calculated: *in vivo* (IV) and *post-mortem* (PM). SNR is expressed in dB (20*ln(RMS(IV)/RMS(PM))) and interpolated in 20 points logarithmically spaced between 10 Hz and 1 kHz.

#### Statistical analysis

Epicortical neural data presented in **Figure 3** are taken from individual measurements on a single animal. In **Figure 3c**, data from 64 electrodes is presented. In **Figure 3d**, data from two selected electrodes is presented. In **Fig. S12a**, impedance data is presented as a histogram with n = 64. The presented impedance at 1 kHz is mean ± SD. In **Fig. SI 12c-d**, the PSD and SNR are calculated from 64 EGNITE electrodes and shown as mean ± SD. In **Fig. SI 12 b**, boxplots of the impedance at 1 kHz for EGNITE (n = 64) and Pt (n = 60) are shown. PSD and SNR from **Fig. SI 12c-d** are calculated from 64 EGNITE electrode and 60 Pt electrodes. For clarity, only the mean is presented.

### Neural Recording: intracortical implantation

Animals were anesthetized induced with a mixture of ketamine/xylazine (75:1, 0.35 ml/28 g i.p.) and anesthesia this state was maintained with an inhalation mask at 1.5% isoflurane. Several micro-screws were placed into the skull to stabilize the implant, and the one on top of the cerebellum was used as a general ground. The probe was implanted in the prefrontal cortex (coordinates: AP: 1.5 mm; ML: ± 0.5 mm; DV: -1.7 mm from bregma). The implantation was performed by using a maltose coating the probe with maltose (see protocol below) to provide temporary probe stiffness and facilitate probe the insertion. The probe was sealed with dental cement. TDT-ZifClip connectors were used to connect the probe to the electrophysiological system via a miniaturized cable (**Figure 3e**). After the surgery, the mouse underwent a recovery period of one week receiving analgesia (buprenorphine) and anti-inflammatory (meloxicam) treatments. Neural activity was recorded with the multi-channel Open Ephys system at a sampling rate of 30 kHz with an Intan RHD2132 amplifier. The experiments of the auditory task were conducted in a soundproofed box, with two speakers inside using protocols based on previously described work.^48^ The sound stimulus consisted of a 15ms-long white noise click, repeated 100 trials (cycles), each separated by 5s (interstimulus interval). During the task, the animal was freely moving.

#### Maltose stiffener protocol

An aqueous solution of maltose is heated up to the glass transition point (Tg), between 130 and 160ºC using a hot plate or a microwave. Once the maltose is viscous, the backside of the probe is brought into contact only with the maltose. As the maltose cools down, it rigidifies and stiffens the probe.

#### Data analysis

Neural signals from each electrode were filtered offline to extract single-unit activity (SUA) and local field potentials (LFPs). SUA was estimated by filtering the signal between 450-6000 Hz and the spikes from individual neurons were sorted using principal component analysis with Offline Sorter v4 (Plexon Inc). To obtain LFPs, signals were downsampled to 1 kHz, detrended and notch-filtered to remove noise line artifacts (50 Hz and its harmonics) with custom-written scripts in Python. **Statistical analysis**. Data shown in Figure 3f are mean ± SD, n = 30 as the number of averaged trials. Data recorded from same electrode is shown at days 28, 58 and 90. Data from a single animal is presented.

### Chronic Epicortical Biocompatibility

#### Surgical implantation of devices

A total of 27 adult, male, Sprague-Dawley rats were used for this study (Charles River, England). Animals were housed at anambient temperature of 21+/-2 ^°^C and humidity of 40-50%, on a 12 h light, 12 h dark cycle. Rats were housed in groups and given free access to diet and water throughout the experimental period. Experimental procedures were carried out in accordance with the Animal Welfare Act (1998), under the approval of the Home Office and local animal welfare ethical review body (AWERB). Animals were anaesthetised with Isoflurane (2-3%) for the duration of surgery, and depth of anaesthesia was monitored using the toe pinch reflex test. Animals were placed in a stereotaxic frame (Kopf, model 900LS), located above a thermal blanket to maintain body temperature. A craniotomy hole (∼ 5 × 4 mm) was made 1 mm away from the midline using a dental drill with a 0.9 mm burr drill bit, the dura was removed and the epicortical device placed on the cortical surface of the brain. The craniotomy hole was sealed with Kwik-sil, followed by dental cement to secure, and the skin sutured closed. Subcutaneous injections of saline (1 mL/kg) and buprenorphine (0.03 mg/kg) were given to replace lost fluids and reduce post-operative pain, and anaesthesia was withdrawn.

#### Tissue collection and processing

Animals were culled at 2, 6 or 12 weeks post-implantation by an appropriate method for the type of analysis to be performed.

#### Histology & Immunohistochemistry

At 2, 6 or 12 weeks post-implantation rats were culled via cardiac perfusion with heparinised (10U/mL, Sigma-Aldrich) phosphate-buffered saline (PBS), followed by 4% paraformaldehyde (PFA, Sigma Aldrich) in PBS. Brains were post-fixed in 4% PFA for 24 hours, then transferred to 30% sucrose in PBS for at least 48 hours before freezing in isopentane. The brains were then stored at -80°C until cryosectioned at 25 μm. The tissue was then stained for ionised calcium binding adaptor molecule 1 (Iba-1) to determine the level of microglial activation. Briefly, tissue sections were blocked with 5% goat serum in PBS with 0.1% Triton-X for 1 hour before overnight incubation at 4 °C with the primary antibody anti-Iba-1 (1:1000, 019-19741, Wako). Sections were then stained with secondary antibody, anti-rabbit Alexa Fluor 594 (A-11012, 1:400; Thermofisher) for 1 hour at room temperature. Slides were mounted with coverslips using Prolong Gold anti-fade mounting media with DAPI (Thermofisher). The probe covered an area of 3 × 3.7 mm on the cortical surface of the brain, tissue sections selected for staining covered 3.2 mm in length of this region. Slides were imaged using the 3D-Histech Pannoramic-250 microscope slide-scanner at 20x and images analysed using CaseViewer (Version 2.4, 3D Histech). To assess for microglia activation, a 3.2 mm area was covered, with one image analysed every 100 μm. Images were taken at 8.5x magnification which detailed a section of the epicortical probe site, 3 mm from the midline of the brain to encompass the area directly under the probe site.

#### Image processing

Microglial activation was analysed using a custom CellProfiler* (Broad Institute, version 3.1.9 from https://cellprofiler.org/) pipeline. First, the EnhanceOrSuppressFeatures module was used to enhance features of the type neurites by applying the tubeness enhancement method. From the enhanced images, cells were segmented using the IdentifyPrimaryObjects module. Preliminary measurements of the cells suggested that the appropriate object diameter range was 3-40 pixels. Objects outside this diameter range or touching the edge of the image were discarded. The cells were segmented using a two class Otsu adaptive thresholding strategy with an adaptive window size of 50 pixels. The objects identified by the IdentifyPrimaryObjects module were input to the MeasureObjectSizeShape module to calculate the necessary properties for cell classification. In the ClassifyObjects module, the category on which to base classifications was specified to be AreaShape, and Extent was selected as the corresponding measurement. The cells were classified as *‘activated’* or *‘non-activated’* based on their Extent property, which is the ratio of the area occupied by the cell to the area occupied by its bounding box. This classification approach was rationalised by the fact that activated microglia have large cell bodies and no processes, and thus occupy a far larger proportion of their bounding boxes than their non-activated counterparts. Finally, the CalculateMath and ExportToSpreadsheet modules were used to calculate and output the desired statistics.

#### Statistical analysis

Data sets are *n* = 3 for each device type: Polyimide-only implant (PI); polyimide with exposed microfabricated gold (gold); and polyimide with microfabricated gold and EGNITE (EGNITE) at all timepoints, with the exception of 6-week GOLD which is *n* = 2 for ELISA data. Contralateral hemispheres were combined at each timepoint to give *n* = 9 at 2 and 12 weeks and *n* = 8 at 6 weeks post-implantation. Graphical representation and analysis of the data was generated using GraphPad Prism software (Version 8). Statistical analysis was completed using a Two-way ANOVA with Tukey’s multiple comparisons test where appropriate; a P value < 0.05 was deemed to be significant.

#### Enzyme-linked Immunosorbent Assay (ELISA)

Following the implantation period, animals were culled by cervical dislocation. Brain tissue was extracted from both the right and left hemisphere of the brain, snap frozen in liquid nitrogen and stored at -80 °C until further use. Tissue was lysed using NP-40 lysis buffer (150 mM NaCl, 50 mM Tric-Cl, 1% Nonidet P40 substitute, Fluka, pH adjusted to 7.4) containing protease and phosphatase inhibitor (Halt™ Protease and Phosphatase Inhibitor Cocktail; Thermofisher Scientific), followed by mechanical disruption of the tissue (TissueLyser LT; Qiagen). Samples were then centrifuged for 10 minutes at 5000 RPM, and the supernatant stored at 4°C until further use. The LEGENDplex™ Rat Inflammation Panel (Cat. No. 740401, BioLegend), a bead-based multiplex ELISA kit was run to quantify the following cytokines; IL-1α, IL-1β, IL-6, IL-10, IL-12p70, IL-17A, IL-18, IL-33, CXCL1 (KC), CCL2 (MCP-1), GM-CSF, IFN-γ and TNF-α. The kit was run according to the manufacturer’s instructions, with protein loaded at a fixed volume of 15 μL. Following incubation with supernatant the beads were run on the BD FACSVerse flow cytometer, and the data analysed using LEGENDplex™ data analysis software.

### Neural Stimulation: intrafascicular implantation

All animal experiments were approved by the Ethical Committee of the Universitat Autònoma de Barcelona in accordance with the European Communities Council Directive 2010/63/EU. Animals were housed at 22±2 °C under a 12:12 h light cycle with food and water ad libitum. The sciatic nerve of anesthetized female Sprague-Dawley rats (250-300 g, ∼18 weeks old) was surgically exposed and the nerve electrodes were implanted transversally across the sciatic nerve with the help of a straight needle attached to a 10-0 loop thread.40 The process was monitored under a dissection microscope to ensure the correct position of the active sites inside the nerve fascicles (**Figure 4b**). During the experiments, the animal body temperature was maintained with a heating pad.

Nerve stimulation was performed by applying trains of biphasic current pulses of a fixed duration of 100 μs and increasing amplitude from 0 to 100 μA in 1 μA steps (Stimulator DS4, Digitimer) through the different EGNITE microelectrodes. Simultaneously, the CMAPs were recorded from GM, TA and PL muscles using small needle electrodes(13 mm long, 0.4 mm Ø, stainless steel needle electrodes A-03-14BEP Bionic) placed in each muscle.^49^ The active electrode was placed on the muscle belly and the reference at the level of the tendon. Electromyography (EMG) recordings were amplified (X100 for GM and TA, X1000 for PL; P511AC amplifiers, Grass), band-pass filtered (3 Hz to 3 kHz) and digitized with a PowerLab recording system (PowerLab16SP, ADInstruments) at 20 kHz.

#### Data analysis

The amplitude of each CMAP was measured from baseline to the maximum negative peak. The voltage peak measurements were normalized to the maximum CMAP amplitude obtained for each muscle in the experiment. A selectivity index (SI) was calculated as the ratio between the normalized CMAP amplitude for one muscle, CMAPi, and the sum of the normalized CMAP amplitudes in the three muscles, following the formula SI_i_ = nCMAP_i_/ΣnCMAP_j_.

### Chronic intraneural biocompatibility

Following a previously reported procedure,^44,5°^ the sciatic nerve of anesthetized Sprague-Dawley female rats (250-300 g, ∼18 weeks old) was exposed and the devices for in vivo biocompatibility with and without EGNITE were longitudinally implanted in the tibial branch of the sciatic nerve (n = 6-8 per group). Briefly, the nerve is pierced at the trifurcation with a straight needle attached to a 10–0 loop thread (STC-6, Ethicon); the thread pulls the arrow-shaped tip of the bent electrode strip. The tip is cut to take away the thread, and the tips of each arm are slightly bent to avoid withdrawal of the device. A longitudinal implant was chosen because it allows a better study of the foreign body response inside the nerve.^44^

#### Nerve and animal functional assessment

Animals were evaluated during follow-up post-implant by means of nerve conduction, algesimetry and walking track locomotion tests.^49^ For conduction tests, the sciatic nerve of the implanted and contralateral paws was stimulated by needle electrodes at the sciatic notch and the compound muscle action potential (CMAP) of the PL muscle was recorded as above. The latency and the amplitude of the CMAP were measured. For the algesimetry test, rats were placed on a wire net platform and a mechanical non-noxious stimulus was applied with a metal tip connected to an electronic Von Frey algesimeter (Bioseb, Chaville, France). The nociceptive threshold (force in grams at which the animals withdrew the paw) of implanted vs contralateral paws was measured. For the walking track test, the plantar surface of the hindpaws was painted with black ink and each rat was left to walk along a corridor. The footprints were collected, and the Sciatic Functional Index calculated.^49^

#### Histology

After 2 or 8 weeks, animals were perfused with PFA (4%), sciatic nerves harvested, postfixed, cryopreserved, and processed for histological analysis. For the evaluation of the foreign body reaction (FBR), sciatic nerves were cut in 15 μm thick transverse sections with a cryostat (Leica CM190). Samples were stained with primary antibodies for myelinated axons (RT97 to label Neurofilament 200K, 1:200, Developmental Studies Hybridoma Bank) and macrophages (iba1, 1:500, Wako). Then, sections were incubated for 1h at room temperature with secondary antibodies donkey anti-Mouse Alexa fluor 488 and donkey anti-Rabbit Alexa fluor 555 (Invitrogen, 1:200). Representative sections from the central part of the implant in the tibial nerve were selected, images taken with an epifluorescence microscope (Eclipse Ni, Nikon) attached to a digital camera (DS-Ri2, Nikon), and image analysis performed with ImageJ software (National Institutes of Health, USA). The amount of Iba1 positive cells in the whole area of the tibial nerve was quantified and the thickness of the tissue capsule was measured as the mean distance of each side of the implant to the closest axons.

#### Statistical analysis

For statistical analysis of data, we used one or two-way ANOVA followed by Bonferroni post hoc test for differences between groups or times. GraphPad Prism software was used for graphical representation and analysis. Statistical significance was considered when p < 0.05.

## Code availability

Custom code developed for neurophysiological analysis is available at https://github.com/aguimera/PhyREC.

## Data availability

The experimental data that support the figures within this paper and other findings of this study can be accessed by contacting the corresponding author. Authors can make data available on request, agreeing on data formats needed.

